# Overexpression of GITRL by B cell IgD low (BDL) B cells is a therapeutic strategy to increase endogenous CD4^+^Foxp3^+^ T regulatory cells for the treatment of autoimmunity

**DOI:** 10.1101/2025.02.04.636512

**Authors:** Mohamed I. Khalil, Cody J. Gurski, Robert Burns, Ryan Zander, Kelli C. Sommers, Angela K. Beltrame, Bonnie N. Dittel

**Author notes:** These authors share first author positions (Mohamed I. Khalil, Cody J. Gurski). All authors have seen and approved the manuscript hasn’t been accepted or published elsewhere. The authors have no competing interests to declare. **Correspondence to**: Bonnie Dittel, Versiti Blood Research Institute, P.O. Box 2178, Milwaukee, WI 53201-2178.

## Abstract

Autoimmune diseases, such as multiple sclerosis (MS), are often chronic with no cures. An underlying commonality of autoimmune diseases is immune-mediated inflammation. Control of inflammation is achieved by steroids and disease-modifying therapies, which can result in severe side-effects. CD4^+^Foxp3^+^ T regulatory cells (Treg), are essential to controlling autoimmune responses and are considered a strong therapeutic target with minimal side effects. To that end, we leveraged our identification of B cell IgD low (BD_L_) B cells that control Treg homeostatic levels in the mouse spleen in a GITRL-dependent manner to demonstrate that overexpression of GITRL by BD_L_ using a B cell-specific GITRL transgene (tg) was sufficient to increase endogenous Treg numbers and attenuate the disease severity of experimental autoimmune encephalomyelitis (EAE), the mouse model of MS. To determine whether increased GITRL expression by BD_L_ could be a therapeutic strategy, WT mice were transplanted with bone marrow from GITRLtg mice. After reconstitution, GITRL expression was increased on BD_L_, Treg numbers were significantly elevated, and EAE was dramatically attenuated. These cumulative data further demonstrate that GITRL is a functional receptor on BD_L_ and its overexpression in B cells is a therapeutic strategy to increase endogenous Treg numbers for treating autoimmunity.

## Introduction

The central nervous system autoimmune disease multiple sclerosis (MS) can be a life-long illness that even when treated with disease-modifying therapies can progress to a stage that is no longer responsive to drugs targeting the immune system (1, 2). Thus, new therapeutic strategies to halt disease progression are greatly needed. To that end, CD4^+^Foxp3^+^ T regulatory cells (Treg) are considered a strong candidate due to their potent suppression of autoimmune responses (3). Humans born with mutations in *FOXP3* lack functional Treg and quickly succumb to lethal IPEX without a bone marrow transplant (4). Similar results were obtained in mice (scurfy) (5). The level of Treg immune suppression is directly correlated to their cell numbers as evidenced by the adoptive transfer of Treg into mice leading to disease suppression in various models of autoimmunity, including the MS model experimental autoimmune encephalomyelitis (EAE) (6, 7). Strategies to increase Treg numbers in autoimmune patients include adoptive transfer of in vitro manipulated Treg and induction and/or expansion of endogenous Treg (3, 8). While early clinical trials have shown safety and some clinical efficacy (9–11), there are currently no FDA-approved Treg-based therapies are on the market.

To develop a novel strategy to increase endogenous Treg numbers, we leveraged our discovery of B cell IgD low (BD_L_) B cells in the mouse that promote homeostatic proliferation of Treg (12, 13). BD_L_ reside in the spleen and emerge from transitional 2 (T2) B cells along the follicular (FO) B cell pathway but are phenotypically and transcriptionally distinct from FO B cells (13). BD_L_ were discovered in a series of studies investigating the mechanism of why mice deficient in B cells due to the loss of IgM heavy chain expression (μMT) did not recover from the signs of EAE (14, 15). μMT mice were shown to have ∼50% reduction in splenic Treg, which were restored by adoptive transfer of BD_L_, but not FO or MZ B cells (12, 13). BD_L_ drive

Treg expansion via engagement of glucocorticoid-induced tumor necrosis factor receptor-related protein ligand (GITRL) with GITR constitutively expressed on Treg (12, 13). Blocking of GITRL on BD_L_ before adoptive transfer blocked Treg expansion and EAE recovery (12, 13). In addition, transgenic (tg) mice overexpressing GITRL (GITRLtg) specifically in B cells were shown to have increased numbers of Treg and attenuated EAE (16), findings we confirmed here.

The above studies suggest that overexpression of GITRL on BD_L_ could be a therapeutic strategy to increase endogenous numbers of Treg. To that end, we first demonstrated that BD_L_ require GITRL expression to attenuate EAE after adoptive transfer. We then utilized single-cell RNA sequencing (scRNAseq) to further refine the BD_L_ phenotype to include a CD24^low^ subset and a CD24^hi^ subset, with the latter displaying a potent capacity to promote Treg expansion and disease recovery following EAE. GITRLtg BD_L_ were superior to WT BD_L_ in inducing Treg expansion after adoptive transfer, but not to sufficient levels to attenuate EAE compared to WT BD_L_. Of importance, GITRL overexpression was not sufficient to endow FO or MZ B cells with the capacity to induce Treg expansion. Next, we asked whether transplantation (T) of WT mice with GITRLtg bone marrow (BM) would recapitulate EAE findings in GITRLtg mice. EAE was strikingly attenuated and accompanied by a significant increase in the absolute number of Treg and their percentage within the CD4 T cell subset as compared to mice transplanted with WT BM. These collective findings, demonstrate that over-expression of GITRL by BD_L_ using BMT is a therapeutic strategy to increase endogenous Treg numbers to attenuate autoimmunity.

## Results

### BD_L_ requires GITRL expression to promote disease recovery and Treg expansion in EAE

In previous studies, we showed that BD_L_ required expression of GITRL to drive Treg expansion in μMT mice (12, 13). Here, those studies were expanded to show that BD_L_ GITRL expression is also required for attenuation of EAE (Fig. 1A, B). BD_L_ were FACS purified and stained with an anti-GITRL blocking antibody before adoptive transfer into μMT mice. EAE was induced by adoptive transfer of myelin basic protein (MBP)-TCRtg encephalitogenic T cells three days later (17). Consistent with our previous studies, WT mice and μMT mice that were adoptively transferred BD_L_ recovered from the signs of EAE, but μMT mice did not recover and exhibited chronic disease (Fig. 1A). Similarly, μMT mice adoptively transferred antibody-blocked BD_L_ did not recover from EAE (Fig. 1A) and exhibited a statistically significant more severe EAE disease as compared to WT and μMT mice transferred BD_L_ when comparing area under the curve (AUC) (Fig. 1B). Treg numbers were quantitated in the spleen at EAE day 30 by gating on CD4^+^Foxp3^+^ Treg. Consistent with our previous studies, μMT mice were found to have a significant reduction in the absolute number of Treg as compared to WT mice, which were significantly increased following transfer of BD_L_ (Fig. 1C). μMT mice that were transferred anti-GITRL blocked BD_L_ did not undergo Treg expansion (Fig. 1C). CD4^+^ T cells (Foxp3^-^) were reduced in all conditions as compared to WT mice with the lowest number found in the GITRL blocking cohort (Fig. 1D). Representative flow cytometry gating for BD_L_ and Treg are shown in Suppl. Fig. 1. These data demonstrate that the primary effector molecule utilized by BD_L_ to promote Treg expansion and EAE recovery is GITRL.

**Figure 1.**
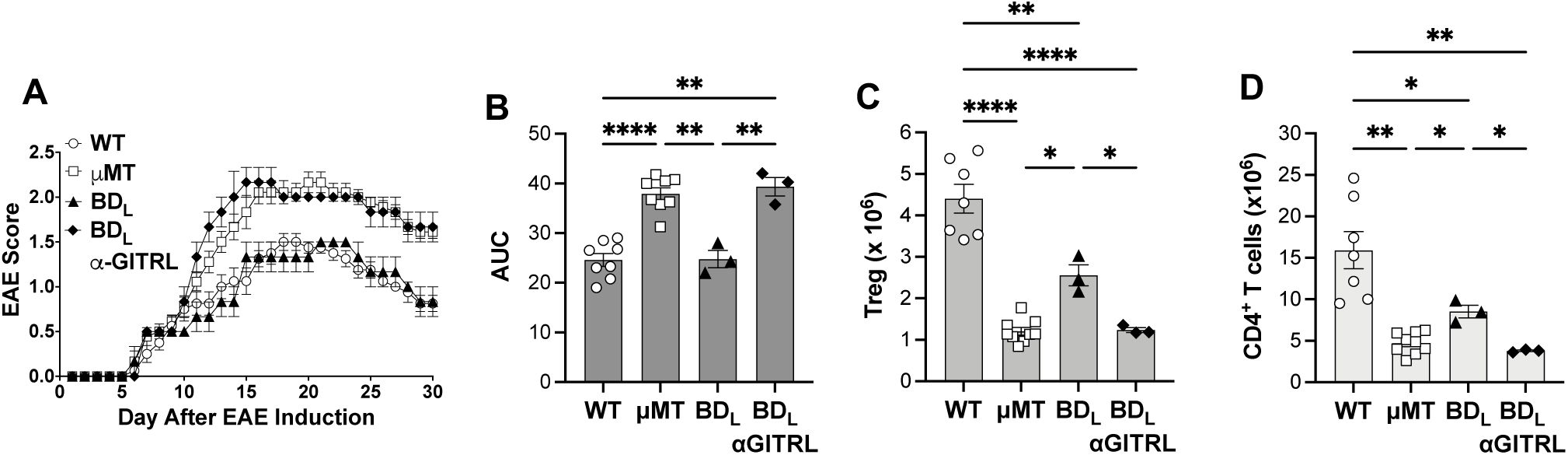
BD_L_ utilize GITRL to attenuate EAE and drive Treg expansion. WT and μMT mice were sublethally irradiated on day −4 and μMT mice were adoptively transferred 4 x 10^6^ FACS purified splenic BD_L_ with or without anti-GITRL blocking on day −3. On day 0 EAE was induced by the adoptive transfer of 1 x 10^6^ MBP-TCRtg encephalitogenic CD4 T cells. Clinical signs of EAE were assessed daily and the average daily disease score (A) and AUC (B) are shown. On EAE day 30 the absolute number of Treg (C) and CD4^+^ T cells (D) were quantitated by flow cytometry. Data shown are the mean ± SEM and each data point represents one mouse from three individual experiments. *p≤0.05; **p<0.01; ***p<0.001; ****p<0.0001. Only significance is shown.

### scRNA-seq Profiling Divides BD_L_ Into Two Major Clusters

Previously, we showed that BD_L_ were functionally and transcriptionally distinct from the mature splenic MZ and FO B cell subsets (12). However, developmental and adoptive transfer studies suggested that the current BD_L_ cell surface phenotype consisted of at least two distinct subsets (12). To address that question, scRNA-seq was performed comparing FACS purified T2, BD_L_, and FO B cells using the 10X Genomics Chromium platform. When monocle was used to perform trajectory analysis the three B cell subsets clustered individually and formed a continuum from T2 to BD_L_ to FO (Fig. 2A). Subsequent UMAP analysis on the BD_L_ subset revealed two major subclusters (subclusters 0 and 1) (Fig. 2B) and one minor subcluster (subcluster 2) that was not further analyzed because it exhibited a unique transcriptional profile that was not consistent with splenic B cell development or naive mature FO B cells (data shown). To identify a cell surface marker that could distinguish between BD_L_ subclusters 0 and 1 a dot plot was generated composed of cell surface receptors that were differentially transcriptionally expressed between the two major subclusters (Fig. 1C). Flow cytometry was used to validate each of the potential markers and found that only CD24 exhibited a distinct bimodal distribution (Suppl. Fig. 3 and Fig. 2D, respectively).

**Figure 2.**
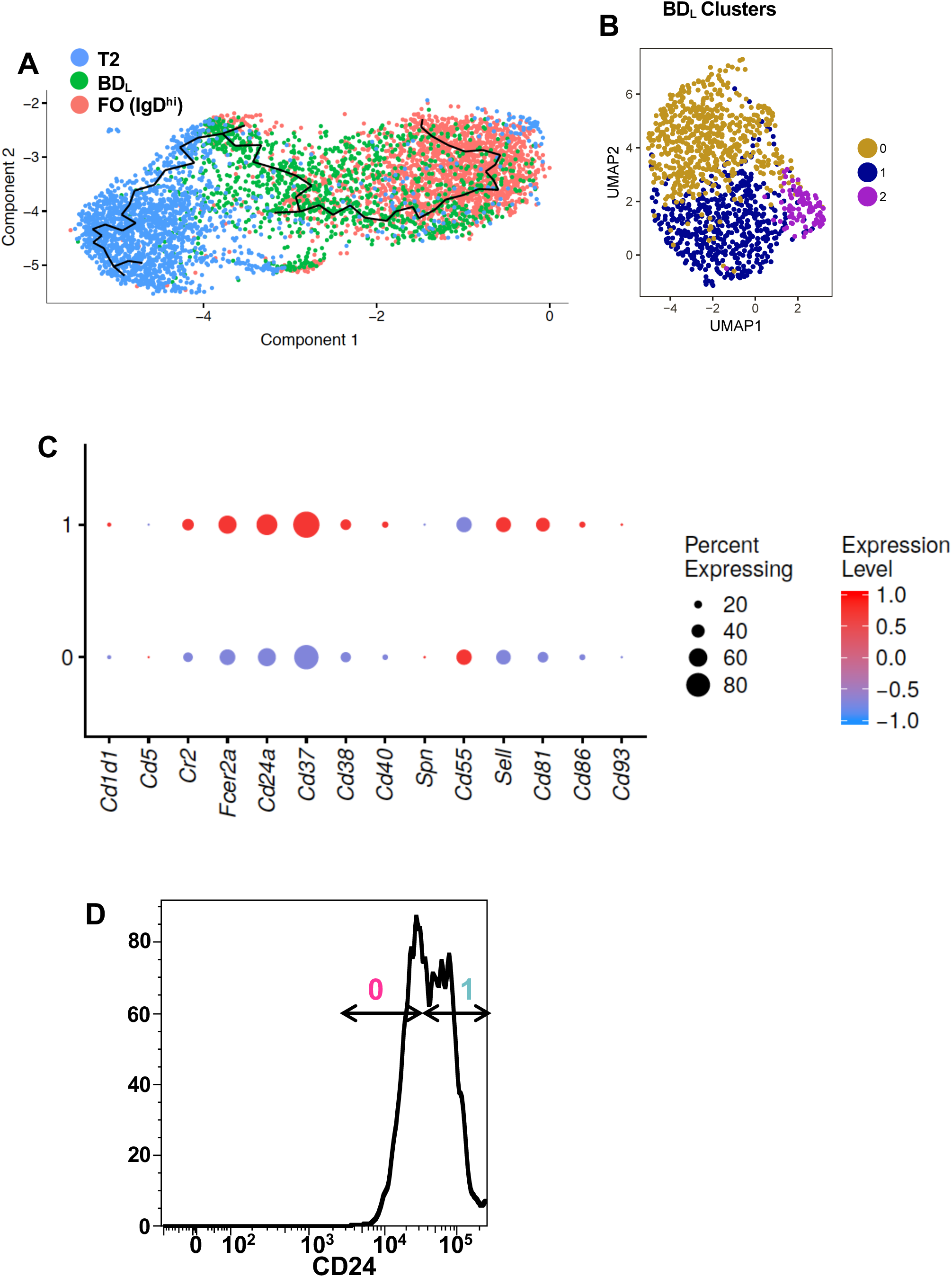
BD_L_ cluster into 2 subclusters that can be separated by CD24 expression levels. Splenic T2, BD_L_ and FO (IgD^hi^) B cells were FACS purified and scRNA-seq was performed and a UMAP was generated including all three subsets overlaid with a trajectory line (A). The BD_L_ subset from A was further clustered revealing three subclusters depicted by UMAP (B). Differential expression of cell surface proteins in subclusters 0 and 1 are shown by dot plots with the size of the dot representing the percent of cells expressing the indicated transcript and expression levels indicated by color with red high and blue low relative expression (C). BDL were stained for CD24 and a flow cytometry histogram was used to indicate a bimodal distribution separating subclusters 0 and 1 (D).

### CD24^hi^ BD_L_ Promote Treg Homeostasis and EAE Recovery

To determine whether CD24 expression could functionally differentiate BD_L_ clusters 0 and 1, BD_L_ were FACS purified separating them into CD24^low^ (subcluster 0) and CD24^hi^ (subcluster 1) subsets (Fig. 2D), adoptively transferred into μMT mice and Treg numbers in the spleen were quantitated seven days later. As we previously reported, μMT mice have a significant reduction in the number of splenic Treg as compared to WT (Fig. 3A) (12, 13). The CD24^low^ subset failed to increase Treg numbers, while the CD24^hi^ subset led to a significant increase in Treg as compared to μMT (Fig. 3A). A similar result was observed for CD4^+^ (Foxp3^-^) conventional T cells (Fig. 3B). Consistent with higher numbers of Treg, both WT mice and μMT mice adoptively transferred CD24^hi^ BD_L_ exhibited EAE recovery, while the CD24^low^ cohort did not recover like μMT mice (Fig. 3C). There was a significant difference in the AUC between the WT and CD24^hi^ groups that recovered and the μMT and CD24^low^ groups that did not (Fig. 3D). The finding that Treg numbers in the spleen were significantly elevated on EAE day 30 in the CD24^hi^ BD_L_ group confirms that they drive and sustain Treg expansion (Fig. 3E). CD4^+^ T cells were also significantly increased after transfer of CD24^hi^ BD_L_ on EAE day 30, but did not reach similar levels as WT mice (Fig. 3F).

**Figure 3.**
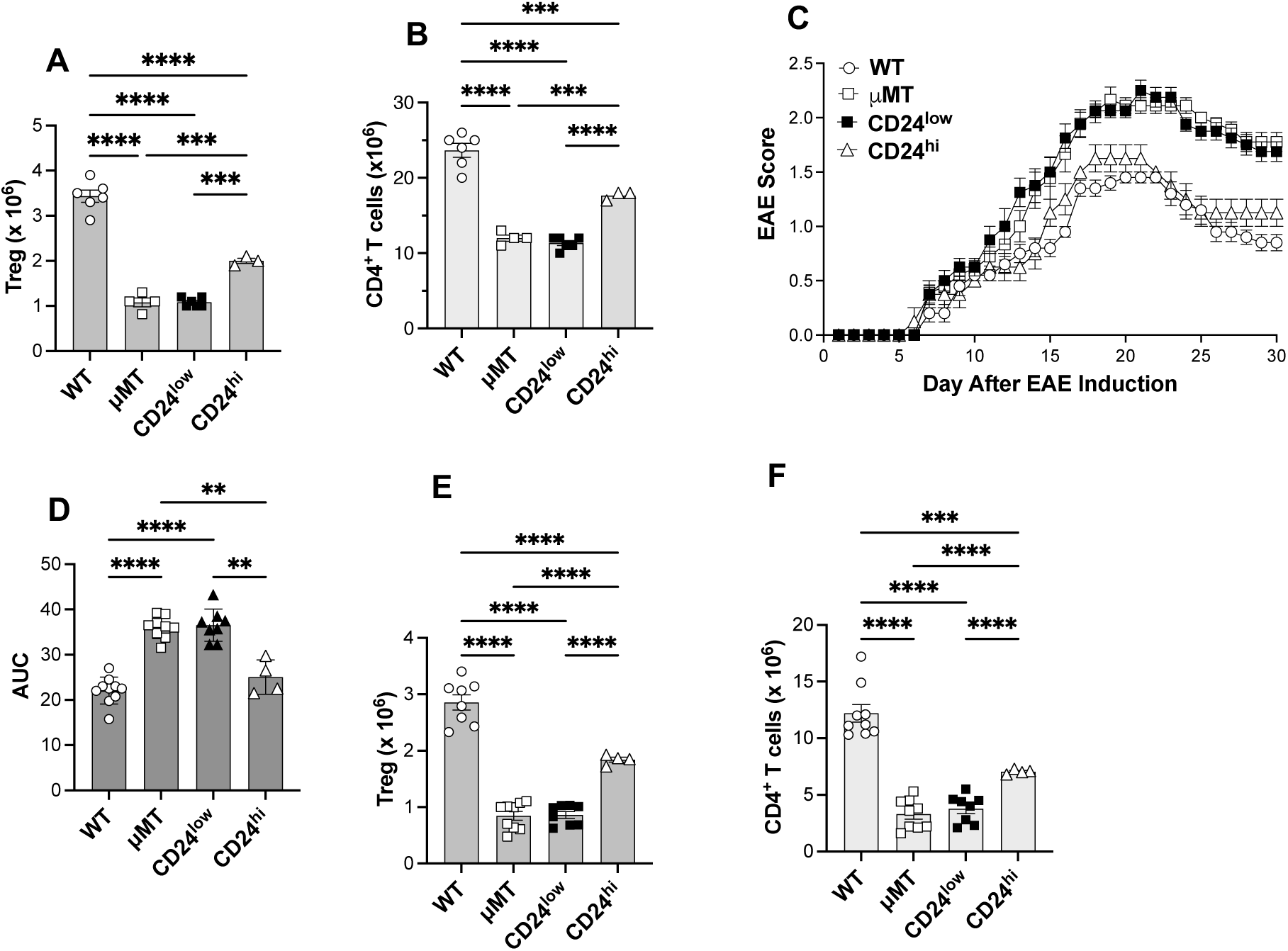
CD24^hi^ BD_L_ exhibit regulatory function, while CD24^low^ do not. μMT mice were adoptively transferred 4-5 x 10^6^ CD24^low^ or CD24^hi^ splenic BD_L_ and 7-10 days later splenic Treg (A) and CD4^+^ T cells (B) were quantitated by flow cytometry. EAE was induced as for Fig. 1A in WT, μMT, and μMT mice were adoptively transferred 4-5 x 10^6^ FACS purified splenic CD24^low^ or CD24^hi^ BD_L_ and average daily disease scores (C) and AUC are shown (D). On EAE day 30 the absolute number of splenic Treg (E) and CD4^+^ T cells (F) were quantitated by flow cytometry. Unmanipulated WT and μMT mice served as controls (A-F). Data shown are the mean ± SEM and each data point represents one mouse from three individual experiments. *p≤0.05; **p<0.01; ***p<0.001; ****p<0.0001. Only significance is shown.

### Characterization of B10.PL GITRLtg mice

To further explore the link between GITRL and Treg expansion, we obtained GITRL B cell-specific transgenic mice (GITRLtg) on the C57BL/6J background (16) and backcrossed them to the B10.PL background. First, it was confirmed that GITRL expression was increased on GITRLtg B cells including MZ, FO, and BD_L_ compared to littermate control mice (Fig. 4A). We also confirmed that B10.PL GITRLtg mice had significant increases in total splenocytes (Fig. 4B), CD4^+^ T cells (Fig. 4C), Treg (Fig. 4D), and percent Treg within the CD4 subset (Fig. 4E) (16). We also confirmed a reported increased proliferation in both Treg and CD4 T cells in GITRLtg mice (data not shown) (16). While increased numbers of splenic B cells were observed in the van Olffen, et. al. study it did not reach significance (16). By increasing the number of mice examined, we found a significant increase in splenic B cells in GITRLtg mice (Fig. 4F) that included marginal zone (MZ) and follicular (FO), but not, BD_L_ B cells (Fig. 4G). To determine whether increased splenic B cell numbers was related to proliferation, Ki-67 expression was measured, with only GITRLtg FO B cells undergoing a significant, but small, increase in the percent of cells proliferating (Fig. 4H). Interestingly, there was no significant difference in the percentage of peripheral blood (PB) CD4^+^ T cells (Fig. 4I) or percent of Treg within the PB CD4 subset in GITRLtg mice (Fig. J).

**Figure 4.**
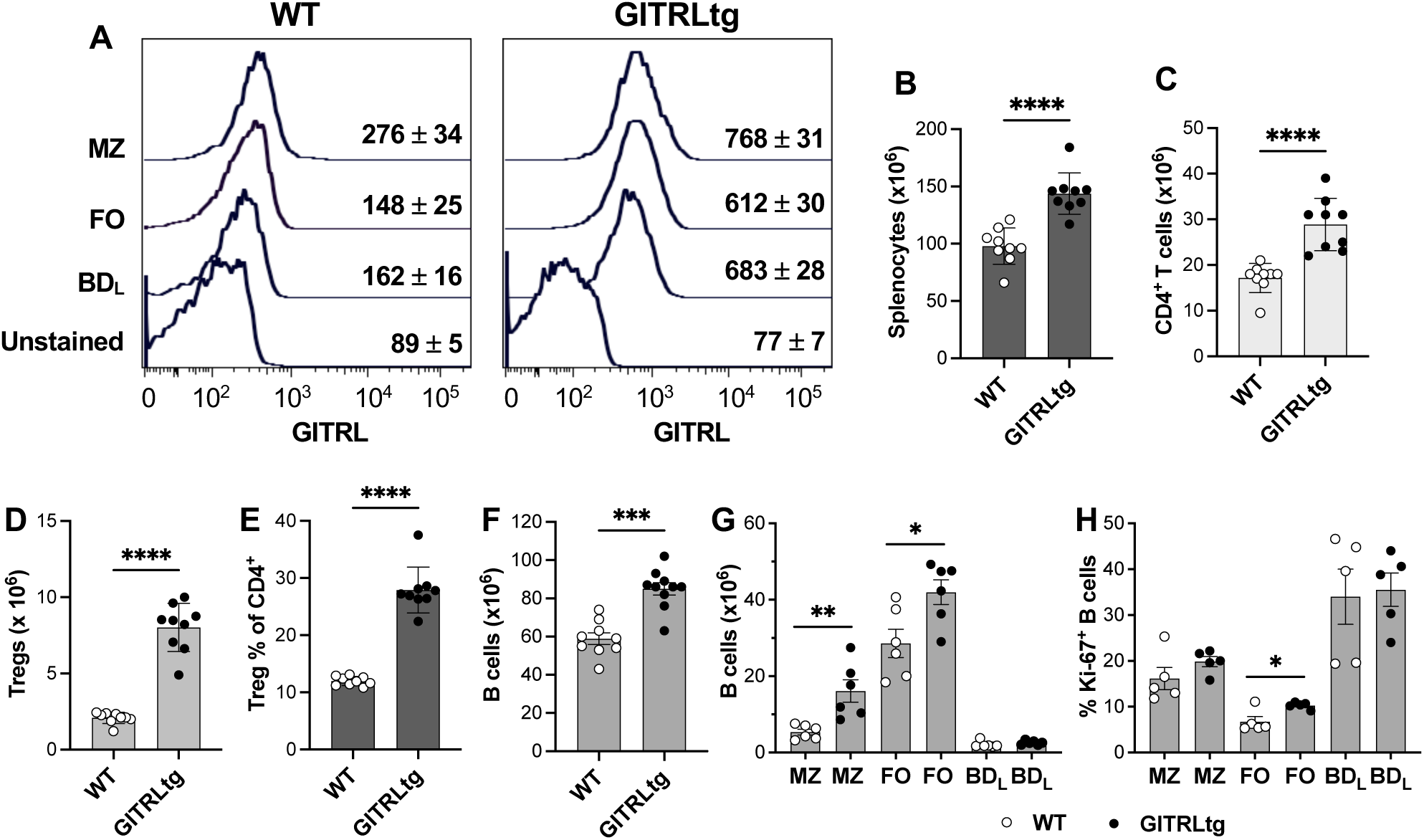
GITRLtg mice have increased cell surface expression of GITRL by B cells and increased numbers of Treg and proportion within CD4^+^ T cells, CD4^+^ T cells, and B cells. Splenic WT and GITRLtg MZ, FO, and BD_L_ B cells were stained for GITRL, and the data are shown as a flow cytometry overlay histogram with the mean fluorescence intensity ± SEM provided for each subset (A). Splenocytes from WT and GITRLtg mice were quantitated by flow cytometry for total splenocytes (B); CD4^+^ T cells (C); Treg (D); percentage of Treg in the CD4^+^ T cell subset (E); CD19^+^ B cells (F); MZ, FO, and BD_L_ B cell subsets (G), and the percent MZ, FO, and BD_L_ B cell subsets expressing Ki-67 (H). Data shown are the mean ± SEM and each data point represents one mouse from two individual experiments. *p≤0.05; **p<0.01; ***p<0.001; ****p<0.0001. Only significance is shown.

### GITRLtg mice exhibit attenuated EAE accompanied by increased numbers of Treg and percentage within the CD4 T cell subset

In previous studies, we positively correlated the number of splenic Treg with recovery from EAE (12, 13). Here that assumption was tested by inducing passive EAE in GITRLtg mice, which have increased numbers of splenic Treg (Fig. 4B) and percentage within the CD4 T cell subset (Fig. 4C). GITRLtg mice succumbed to EAE like WT mice in terms of 100% incidence (data not shown), day of onset, and kinetics of early disease (Fig. 5A). At day 15 the curves split with WT mice continuing to progress and GITRLtg mice plateauing (Fig. 5A). Both groups underwent recovery starting on ∼day 21, with GITRLtg exhibiting less severe EAE as shown by both disease curves (Fig. 5A) and AUC (Fig. 5B). On EAE day 30 both splenic Treg (Fig. 5C) and CD4 T cells (Fig. 5D) were significantly increased in the spleen of GITRLtg mice. In addition, the percent of Treg within the CD4^+^ T cell subset was also significantly increased (Fig. 5E). To determine whether peripheral cell numbers were reflected in the CNS, mononuclear cells were isolated from the brain and spinal cords at peak WT EAE (Day 18) and total CD11b^+^ myeloid cells was used as a marker of overall inflammation. WT mice had significantly higher numbers of myeloid calls as compared to GITRLtg mice, consistent with more severe EAE (Fig. 5G). Both CD4^+^ T cells (Fig. 5G) and Treg (Fig. 5H) were significantly increased, like the spleen. Interestingly, the percent of Treg within the CD4^+^ T cells was not increased in the CNS during EAE (Fig. 5I). These cumulative data provide strong evidence that total Treg numbers are directly correlated to severity of EAE.

**Figure 5.**
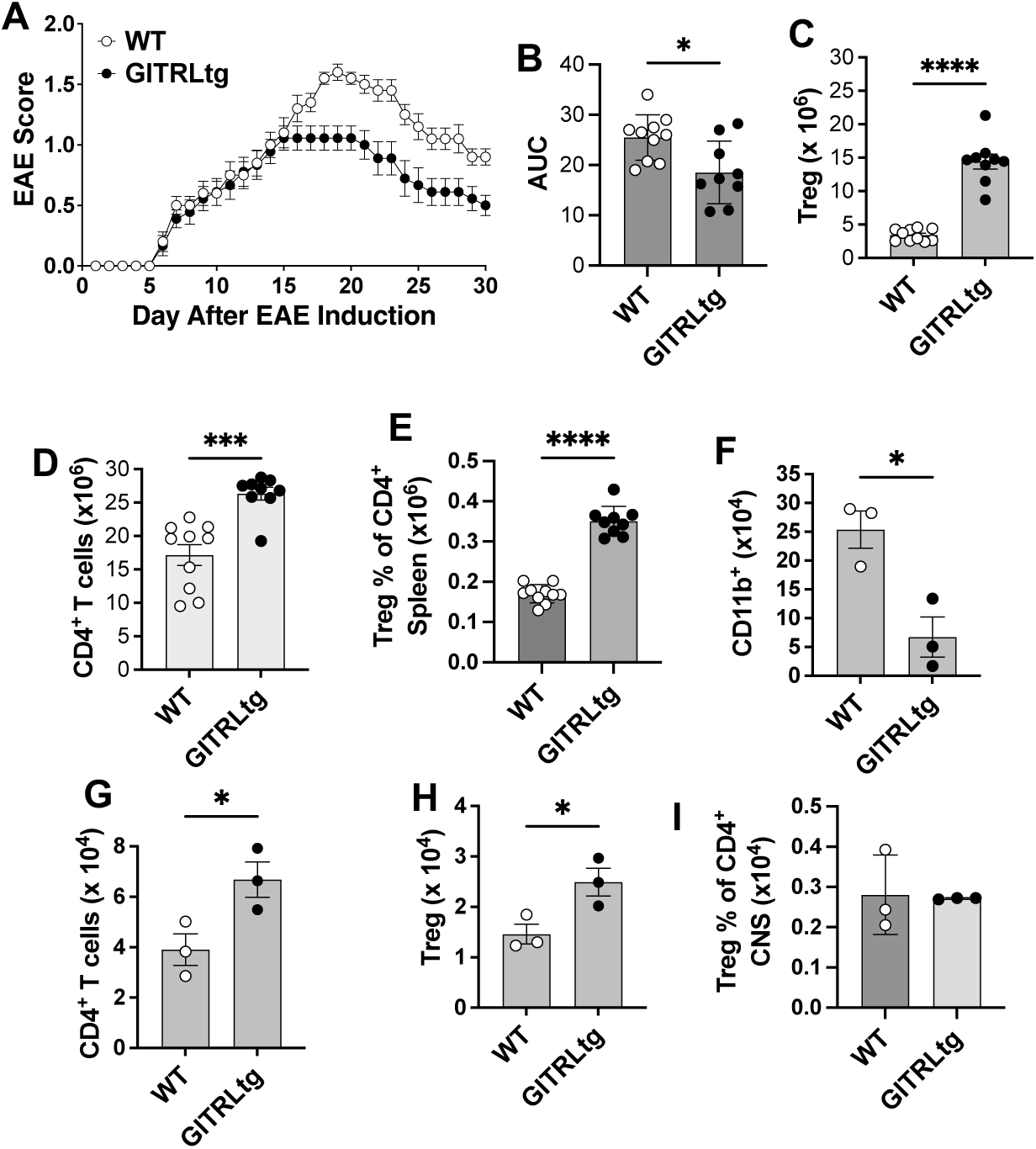
GITRLtg mice exhibited attenuated EAE accompanied by increased numbers and proportions of Treg. EAE was induced as for Fig. 1A in WT and GITRLtg mice and average daily disease scores (A) and AUC are shown (B). On EAE day 30 the absolute number of splenic Treg (C) and CD4^+^ T cells (D) were quantitated by flow cytometry. The percentage of Treg within the CD4 T cell subset was calculated from flow cytometry data (E). On EAE day 18, mononuclear cells were isolated from the brains and spinal cord of WT and GITRLtg mice and the absolute number of CD11b^+^ (F), CD4^+^ T cells (G), Treg (H), and percentage of Treg within the CD4 T cell subset (I) was calculated by flow cytometry. Data shown are the mean ± SEM and each data point represents one mouse from three individual experiments. *p≤0.05; ***p<0.001; ****p<0.0001. Only significance is shown.

### GITRLtg BD_L_ are more efficient at Treg expansion after adoptive transfer as compared to WT BD_L_, but not to a sufficient level to attenuate EAE

Next, the therapeutic potential of increased GITRL expression by BD_L_ was tested. As previously shown, μMT mice have a significant reduction in splenic Treg as compared to WT mice (Fig. 6A). Following the adoptive transfer of WT and GITRLtg BD_L_ into μMT mice, both led to a significant increase in Treg compared to μMT mice, with GITRLtg being significantly more efficacious (Fig. 6A). But, neither BD_L_ population drove Treg numbers to WT levels (Fig. 6A). Neither overexpression of GITRL by FO nor MZ B cells was sufficient to afford them regulatory activity (Fig. 6A). The adoptive transfer was repeated and EAE was induced with more severe disease observed in μMT mice when adoptively transferred GITRLtg FO or GITRLtg MZ B cells (Fig. 6B). In contrast, μMT mice adoptively transferred WT BD_L_ and GITRLtg BD_L_ had a similar disease course as WT mice accompanied by recovery (Fig. 6B). Significance is shown by AUC (Fig. 6C). Treg numbers in the spleen after WT and GITRLtg BD_L_ adoptive transfer were sustained on EAE day 30 (Fig. 6D). In contrast, there was little change in CD4^+^ T cell numbers in all the groups compared to WT mice (Fig. 6E). These data demonstrate that adoptive transfer of BD_L_ that overexpress GITRL allow uMT mice to recover from EAE, but not better than adoptive transfer of WT BD_L_.

**Figure 6.**
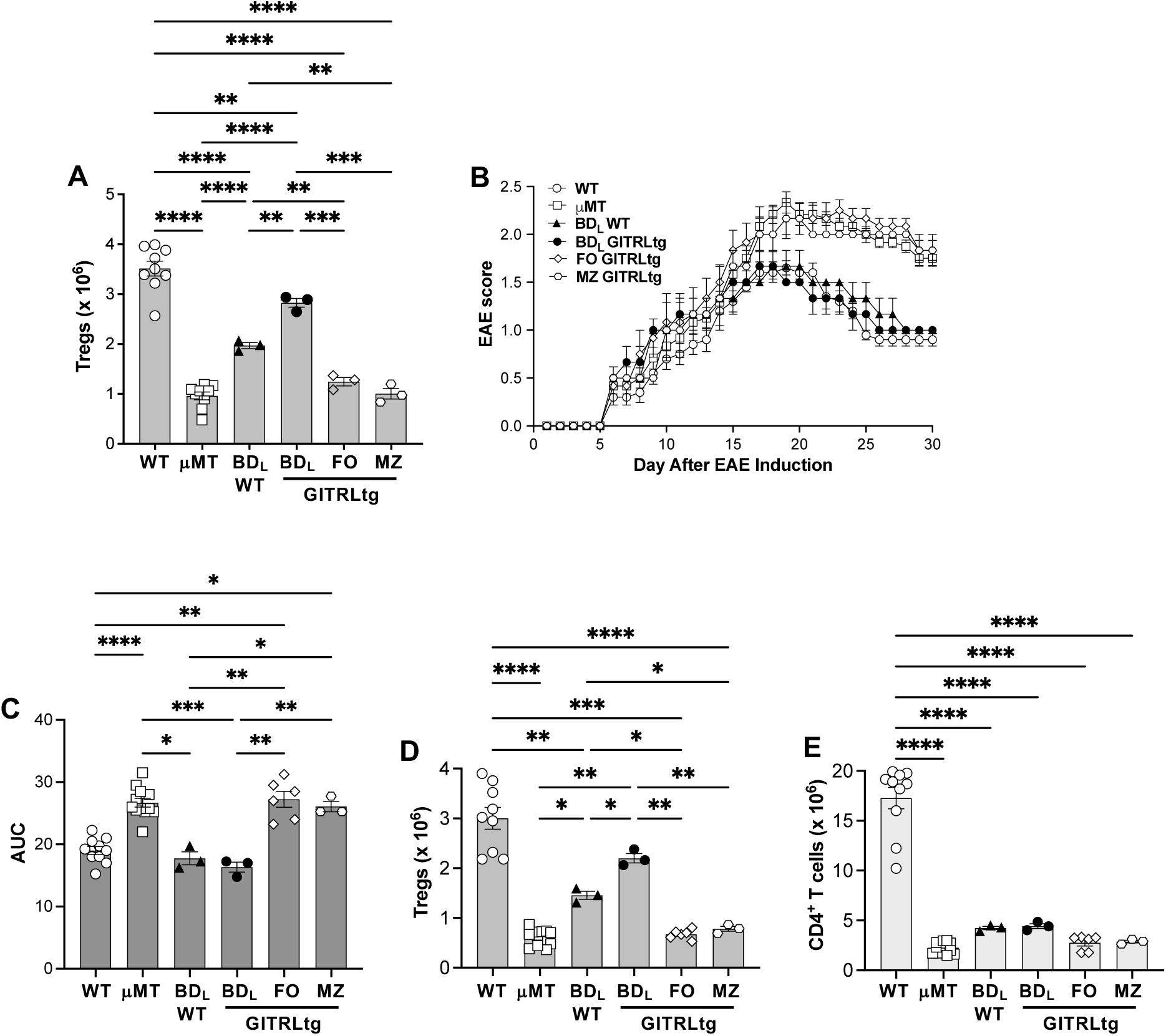
Adoptive transfer of GITRLtg BD_L_ is not sufficient to increase Treg numbers to therapeutic levels. μMT mice were adoptively transferred 4-5 x 10^6^ WT BD_L_, GITRLtg BDL, GITRLtg FO, or GITRLtg MZ splenic B cells. Splenic Treg were quantitated 7-10 days later by flow cytometry (A) or EAE was induced as for Fig. 1A and the average daily disease scores (B) and AUC are shown (C). On EAE day 30 the absolute number of splenic Treg (D) and CD4^+^ T cells (E) were quantitated by flow cytometry. Unmanipulated WT and μMT mice served as controls (A-E). Data shown are the mean ± SEM and each data point represents one mouse from three individual experiments. *p≤0.05; **p<0.01; ***p<0.001; ****p<0.0001. Only significance is shown.

### Transplantation of GITRLtg donor BM into WT mice led to a profound reduction in EAE severity correlated with increased numbers of Treg

Because the adoptive transfer of GITRLtg BD_L_ into μMT mice did not result in sufficient numbers of Treg to attenuate EAE (Fig. 6A, B), an alternative strategy of transplantation of GITRLtg BM into WT recipient mice (GITRLtg→WT) was utilized. Transplantation of WT BM into WT mice (WT→WT) served a control. After 12-14 weeks of reconstitution, EAE was induced by active induction via immunization with the MPB peptide Ac_1-11_. EAE disease was significantly attenuated in mice transplanted with GITRLtg BM, as shown by disease curves (Fig. 7A) and AUC (Fig. 7B). At EAE day 30, WT→WT mice without EAE had comparable numbers of total splenocytes, CD4^+^ T cells, CD19^+^ B cells, and BD_L_ (Fig. 7C-F, respectively) as unmanipulated WT mice (Fig. 4). Interestingly, transplantation of GITRLtg BM into WT mice reset total splenocytes and subsets to WT levels (Fig. 7C-F), as compared to significantly higher cell numbers in unmanipulated GITRLtg mice (Fig. 4). As expected, WT→WT EAE mice exhibited a significant loss in the four subsets (Fig. 7C-F), that was prevented in the GITRLtg→WT cohort (Fig. 7C-F).

**Figure 7.**
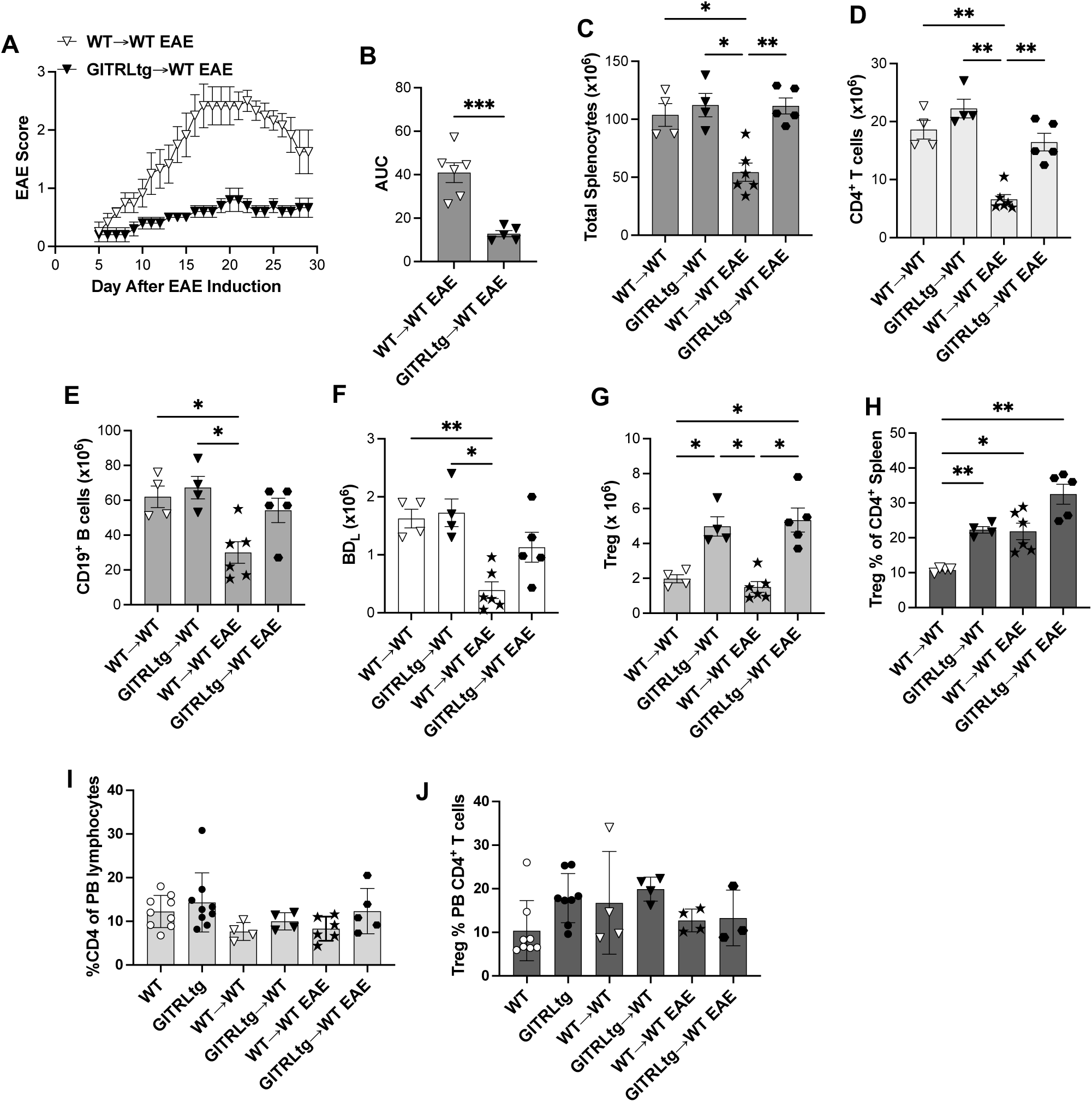
Transplantation of WT mice with GITRLtg BM leads to attenuated EAE associated with increased numbers and proportions of Treg. WT mice were transplanted with WT (WT→WT) or GITRLtg (GITRLtg→WT) BM and after 12-14weeks, EAE was induced by immunization with the MBP Ac_1-11_ peptide and signs of disease was assessed daily starting and the average daily disease scores (A) and AUC are shown (B). On EAE day 30 the absolute number of splenocytes (C), CD4^+^ T cells (D), CD19^+^ B cells (E), BD_L_ (F), Treg (G), percentage of Treg within the CD4 T cell subset (I) was calculated by flow cytometry. In the PB on EAE day 30 the percent CD4^+^ T cells within the PB lymphocyte gate (I) and percentage of Treg in the CD4^+^ T cell subset (J) was quantitated by flow cytometry. Unmanipulated WT and GITRLtg mice served as controls (I, J). In the GITRLtg→WT EAE cohort one mouse was removed from the data set due to insufficient BM reconstitution. Data shown are the mean ± SEM and each data point represents one mouse from three individual experiments. *p≤0.05; **p<0.01; ****p<0.0001. Only significance is shown.

Total splenic Treg numbers were not impacted by EAE (Fig. 7G). Like GITRLtg mice (Fig. 4D) GITRLtg→WT mice with or without EAE had significantly more Treg than the WT→WT cohorts (Fig. 7G). EAE significantly increased the percentage of Treg within the CD4^+^ T cell subset in both WT→WT and GITRLtg→WT cohorts (Fig. 7H), with the latter cohort reaching levels observed in GITRLtg mice (Fig. 4E). One aspect of Treg therapy is whether their increased cell numbers can be tracked in the PB. To address this question, first the percentage of PB CD4^+^ T cells was determined, which was similar among all cohorts of mice (Fig. 7I).

Similarly, the percentage of Treg within the CD4^+^ T cell subset varied little among the six cohorts of mice (Fig. 7J). These collective data indicate that PB CD4^+^ T cell homeostatic proportions are stable in health and disease and do not reflect cellular changes in the spleen. Overall, the therapeutic strategy of BMT with BD_L_ overexpressing GITRL was shown to be effective at reducing signs of EAE.

## Discussion

In this study, we investigated whether over-expression of GITRL on BD_L_ could be a therapeutic strategy to increase endogenous numbers of Treg to attenuate EAE severity. To decrease patient burden, an ideal therapeutic would be a one-time treatment. Using a refined CD24^hi^ BD_L_ phenotype, we showed that adoptive transfer of GITRLtg BD_L_, but not FO or MZ B cells, drove Treg expansion after adoptive transfer into μMT mice. However, the increased numbers of Treg did not reach that of GITRLtg mice and were not sufficient to attenuate EAE as compared to WT BD_L_. These data suggested that one-time adoptive transfer of BD_L_ overexpressing GITRL was not a viable strategy to sustain high levels of Treg. To that end, we reasoned that GITRL overexpression by BDL would need to be sustained, such as in GITRLtg mice. To achieve that goal, WT mice were transplanted with GITRLtg BM leading to increased GITRL expression by B cells, increased numbers and percentages of Treg, most importantly highly attenuated EAE. These cumulative data suggest that genetically increasing GITRL expression by BD_L_ is a therapeutic strategy to drive increased numbers of endogenous Treg numbers and attenuate autoimmunity.

In previous studies, we showed that BD_L_ are phenotypically and transcriptionally related to the FO B cell lineage and emerge from the T2 B cell subset in the spleen (12). To better define the development trajectory of BD_L_ and to find new markers, scRNAseq was utilized. Subclustering revealed that BD_L_ were composed of two major subsets, which could be differentiated by CD24 expression levels (Fig. 2D). Remarkably, the CD24^hi^, but not the CD24^low^ cluster, drove Treg expansion after adoptive transfer into μMT mice and EAE recovery (Fig. 3). It is unclear whether BD_L_ are a unique B cell subset or a developmental stage along the FO pathway. B cell subsets are challenging to phenotype because there is no single marker that can differentiate each developmental stage but relies on differential expression of a collection of markers (18–20). In the spleen the FO lineage develops along a trajectory of immature, T1, T2, and then FO (18–20). Previously we showed that after BMT T2 appeared at day 13 and FO at day 15, with BD_L_ having a slightly earlier emergence than FO B cells (12). This is consistent with pseudotime analysis predicting that BD_L_ lie along the FO pathway between T2 and mature FO B cells (Fig. 2A). Previously, IgD^low/-^ was used as the sole marker differentiating BD_L_ from FO B cells, both of which express intermediate levels of IgM. Here we have identified CD24^hi^ as an additional marker. T2 B cells are phenotyped as IgM^hi^IgD^hi^CD24^hi^ and mature FO B cells as IgM^int^IgD^hi^CD24^low^ (19, 21, 22) while BD_L_ are IgM^int^IgD^low/-^CD24^hi^. While the BDL CD24^hi^ phenotype is consistent with a development path just after T2, the IgD^low/-^ phenotype is more consistent with MZ or T1 B cells (19, 21, 22). While the precise development pathway of BD_L_ is unclear and being investigated in other studies, the finding that only BD_L_ have the capacity to drive Treg expansion demonstrate that they are distinct from FO and MZ B cells. Of clinical importance is the finding that overexpression of GITRL was not sufficient to endow FO and MZ B cells with regulatory activity.

Using B cells therapeutically is challenging because upon adoptive transfer they cannot persist by homeostatic expansion and have a finite lifespan unless they differentiate into plasma cells (23). However, studies have shown that B cells can be used therapeutically in humans (24). An Epstein-Barr virus lymphoblastoid cells line expressing a p21 ras mutation was used in a small clinical trial as antigen presenting cells to drive anti-tumor T cell responses in pancreatic carcinoma patients (25). In patients receiving allogeneic hematopoietic stem cell transplantation, CD19^+^ PB B cells from donors were adoptively transferred after immunosuppression was tapered, which were shown to generate a vaccine-specific response indicating generation of plasma cells (26). Other strategies targeted at tumors have been validated in mice (24). A major roadblock to utilizing B cells for cell therapy is how to genetically modify them for specific functions. Strategies include viral vectors including lentiviral and adeno-associated virus used to transduce primary B cells (27). Here, we demonstrate an alternative strategy of using BMT using genetically modified stem cells (Fig. 7) sourcing BM from GITRLtg with transgene expression restricted to B cells. Translation to humans would require genetic manipulation of stem cells prior to BMT, an approach that has been FDA approved for treating sickle cell disease using lentiviral vectors. In mice, we have demonstrated that lentiviral transduction of Sca1^+^ BM stem cells driving myelin oligodendrocyte glycoprotein in platelets facilitated immune tolerance leading to attenuation of EAE (28). Collectively, these findings provide a proof-of-concept that stem cells could be genetically modified leading to increased expression of GITRL in B cells leading to increased homeostatic levels of Treg and in turn greater immune suppression of autoreactive immune responses.

The primary change in lymphocytes observed in GITRLtg was an increase in the percentage of Treg within the conventional CD4 subset in the spleen (Fig. 4D), but interestingly this was not observed in GITRLtg→WT mice (Fig. 7D). Numerous studies have shown that it is the proportion of Treg more than their absolute cell numbers that determines the extent of their immune suppression. Our data showing that increasing the proportion of Treg via modulation of B cell GITRL expression supports those previous findings. Treg are considered a strong therapeutic target for the treatment of autoimmunity, including MS (29). Current strategies include in vitro expanding patient Treg and adoptively transferring them back to the patients, with clinical trials in diabetes and graft-versus-host disease (GVHD) showing the approach was safe with some clinical benefit (30–32). A major drawback to Treg adoptive cell transfer is the lack of Foxp3 stability leading to conversion to conventional CD4^+^ T cells (33, 34). A second approach involves the use of biologics/small molecules to induce the expansion of Treg in situ, with the caveat that the therapeutic could be immunogenic (35–37). Our BMT strategy of BD_L_ expanding endogenous Treg should last the life of the patient avoiding lifelong treatments. This will reduce both patient morbidity and dramatically reduce the cost of treatment over a lifetime. However, BMT does come with substantial risk including infections and death (38, 39). Some risks can be mitigated with autologous BMT that does not result in GVHD. In MS, autologous hematopoietic stem cell transplantation (aHSCT) has been performed with the goal of resetting the immune system and has shown efficacy (40, 41). A comprehensive review of published studies comparing aHSCT to various DMT concluded that aHSCT was superior in reducing MRI lesions, disability scores, progression, and relapse rates in MS patients (41, 42). Most of the studies did not report long-term follow-up (years), but the limited data showed worsening of disease or relapse in 32-56% of patients (42). While these reports are promising, additional strategies are required to further reduce worsening of disease following aHSCT. By combining aHSCT with BD_L_ GITRL-Treg therapy we propose that progression and relapse rates will be further reduced leading to an important clinical impact on the quality of life for MS patients.

One challenge to our studies is that while Treg proportions within the conventional CD4 T cell subset is increased in the spleen in mice with increased B cell GITRL expression, this is not reflected in the PB (data not shown). Interestingly, a similar finding was observed in the CNS of EAE mice (Fig. 5I). These data suggest that Treg that enter the CNS during EAE are from the PB further indicating that Treg immune suppression in EAE is largely occurring in the spleen or other secondary lymphoid organs. Whether Treg suppression in the CNS plays a major role in controlling CNS inflammation remains unresolved. Regarding Treg homeostatic levels, we have shown that B cell deficient mice due to genetic ablation (μMT) or antibody depletion have an ∼50 reduction in total splenic Treg (12). It is likely that the remaining 50% of Treg are being maintained by dendritic cells in an MHC class II and/or CD80/86-dependent manner (43, 44). This suggests two populations of Treg, under different homeostatic control, one localized to the spleen that interact with BD_L_ and a second circulating subset under dendritic cell control. Whether these two proposed subsets differ in phenotype and function is unknown.

In this study, we further defined the phenotype of BD_L_ to include a CD24^hi^ cell surface phenotype which was shown to partition the original BD_L_ phenotype by ∼50%. Although CD24 is not a functional marker, additional evidence was provided that GITRL is not only a functional marker, but its function can be enhanced by increasing its expression on BD_L_. After adoptive transfer GITRLtg BD_L_ were superior to WT BD_L_ in driving Treg expansion, but not to sufficient numbers to attenuate EAE. Of importance, neither overexpression of GITRL on FO nor MZ B cells afforded them regulatory activity. Because GITRLtg mice have increased numbers of Treg and attenuated EAE, we tested whether a BMT approach could be used as a therapeutic strategy to increase endogenous Treg numbers. To that end, transplantation of GITRLtg BM into WT mice led to a significant increase in Treg numbers and greatly attenuated EAE. These cumulative data provide evidence that overexpression of GITRL by BD_L_ is a viable strategy to increase endogenous Treg numbers to attenuate autoimmunity.

## Materials and Methods

### Mice

B10.PL (B10.PL-H2^u^ H2-T18^a^/(73NS)SnJ) mice were purchased from The Jackson Laboratory and then maintained in the Dittel animal colony (Bar Harbor, ME). GITRLtg mice on the C57BL/6 background were provided by Dr. Martijn Nolte and crossed with B10.PL mice to change the MHC locus to H-2^u^ and maintained as GITRLtg hemizygotes(16). B10.PL MBP-TCRtg were maintained as hemizygotes and B10.PL μMT mice were previously described (14, 17). All mice were housed and bred in the Biomedical Resource Center of Medical College of Wisconsin (MCW) and animal protocols using all relevant ethical regulations approved by the MCW Institutional Animal Care and Use Committee. All experiments were carried out with mice that were age- and sex-matched utilizing both sexes.

### Peptide and antibodies

MBP Ac_1–11_ peptide (Ac-ASQKRPSQRSK) was generated by the Protein Core Laboratory of the Blood Research Institute, Versiti Wisconsin. Commercial antibodies utilized in this study are described in Supplementary Table 1.

### EAE induction

EAE was passively induced by the i.v. adoptive transfer of 1 × 10^6^ MBP-specific encephalitogenic T cells generated in vitro from MBP-TCRtg mice by activation with MBP Ac_1–11_ peptide into sublethally irradiated (360–380 rad) mice as described (17, 45) or by actively induced by subcutaneous immunization with the MPB peptide Ac_1-11_ (300 μg) emulsified in complete Freund’s adjuvant (Condrex, Inc, Woodinville, WA) with pertussis toxin (300 ng) (List Biological Laboratories, Campbell, CA) administered intraperitoneally on days 0 and 2. Clinical signs of EAE were scored daily as follows: 0, no disease; 1, limp tail; 1.5, hind limb ataxia; 2, hind limb paresis; 2.5, partial hind limb paralysis; 3, total hind limb paralysis; 4, hind and fore limb paralysis; and 5, death.

### Single-Cell RNA-seq

All samples were demultiplexed, aligned to the *Mus musculus* mm10 genome, and gene x cell UMI count matrices were generated using 10X Genomics cellranger v2.2.0 pipeline. UMI count matrices were imported into R v3.5.3 as Seurat v3 objects (46). Cells with less than 400 genes detected and greater than 6% mitochondrial transcript content were discarded. Cell types were predicted using the SingleR v1.0 library (47) in conjunction with the Mouse Immunological Genome Project (ImmGen) transcriptome database (48). Based on these cell type predictions all contaminating cells were removed and only B cells were retained for further analysis. For the remaining cells, gene expression values were normalized using the LogNormalization method with a scaling factor of 10,000. Two-thousand variable features were detected for further use in clustering using the variance stabilizing transformation method. All normalized gene expression values were scaled, centered, and principal component analysis was conducted. Clusters were detected using Louvain’s method (49) using the first 10 principal components with a resolution value of 0.5. Clusters were visualized using the uniform manifold approximation and projection method (50), also using the first 15 principal components. Cluster 2 was mainly comprised of IgD low cells, so this cluster was subset and further subclustered to explore IgD low cell heterogeneity. Again, variable features were detected within cluster 2 cells, data were re-scaled, and principal component analysis was run. Cluster 2 cells were re-clustered using the first 5 principal components using a resolution of 0.3. Two subclusters within cluster 2 were identified and differential gene expression was conducted between these two subclusters using the MAST method (51). Monocle v3 was used to order cells from all B cell clusters and sub-clusters in pseudotime (52). VIPER was used to calculate transcription factor activities across all clusters and subclusters using the Dorothea transcription factor target database (53).

### Single cell isolation and flow cytometry

Single-cell suspensions from mouse spleens were obtained by mincing with glass slides followed by RBC lysis. Peripheral blood lymphocytes were collected by submental bleeding mice directly into ACK RBC lysis buffer. Blood was incubated for 15 minutes at room temperature before tubes were washed with FACS buffer and centrifuged at 400xg. Tubes were aspirated and the pellets re-suspended into antibody mixes. Brains and spinal cords from mice intracardially perfused with PBS were homogenized and mononuclear cells were isolated using 40%/70% discontinuous Percoll gradients. FcR were blocked with anti-mouse FcR (2.4G2). Single cell suspensions were counted and up to 1 x 10^6^ cells were stained with specific antibodies. Cells were acquired on an LSRII or Celesta flow cytometers (BD Biosciences, San Diego, CA, USA) and gated for singlets prior to acquisition. Data were analyzed using the FlowJo software (BD Biosciences, San Diego, CA, USA).

### Fluorescence activated cell sorting (FACS) and adoptive transfer of B cells

Total splenic B cells from 8–10-week-old mice were purified by negative selection using magnetic cell sorting (STEMCELL Technologies, Vancouver, BC, Canada) followed by FACS purification of BD_L_ (B220^+^IgM^+^CD21^int^CD23^+^CD93^−^IgD^low/-^), CD24^hi^ BD_L_, CD24^lo^ BD_L_, FO (B220^+^IgM^+^CD21^int^CD23^+^CD93^-^IgD^hi^), and MZ (B220^+^IgM^hi^CD21^hi^CD23^-^CD93^-^) B cells using a FACSAria cell sorter (BD Biosciences, San Diego, CA, USA). B cell purities were ∼99% as determined by flow cytometry. For some experiments FACS-purified Bells were incubated with anti-mouse GITRL (10 μg/ml) at 4° C for 60 min for GITRL blocking. B cells were washed in PBS and 5-20 x 10^6^ cells were i.v. injected into each recipient mouse. EAE was induced three days later. Splenic CD4^+^Foxp3^+^ and CD4^+^ T cells were enumerated on day 7-10 by flow cytometry.

### BMT

Total BM cells (6 x 10^6^) from WT or GITRLtg mice was i.v. injected into lethally irradiated (950R) recipient WT mice and allowed to reconstitute for 12-14 weeks.

### Statistical analysis

Data were analyzed using GraphPad prism (San Diego, CA) and were presented as mean ± SEM. Statistical significance comparing two groups were determined using the unpaired *t* -test and multiple group comparisons were conducted using Welch and Brown-Forsythe one-way ANOVA comparing the mean of each column with the mean of every other column. *P* values ≤0.05 were considered significant.

## Author contributions

BND conceived of the studies and secured funding; MIK, CJG, RZ, and BND designed the research studies; MIK, CJG, RZ, KCS, and AKB conducted experiments; MIK, CJG, KCS, and AKB acquired data; MIK, CJG, RB, KCS, AKB, and BND analyzed data; and all authors assisted with writing the manuscript. First authorship is shared by MIK and CJG, which was determined by critical individual contributions by both to the data.

## Acknowledgements

The authors would like to thank Versiti Blood Research Institute Shared Resources (RRID: SCR_025503) for their services, instrumentation, and specialist support. The authors would like to thank Dr. Martijn Nolte for the generous gift of the GITRLtg mouse (current address ZonMW, The Netherlands) and……

---

**Supplemental Figure 1.**
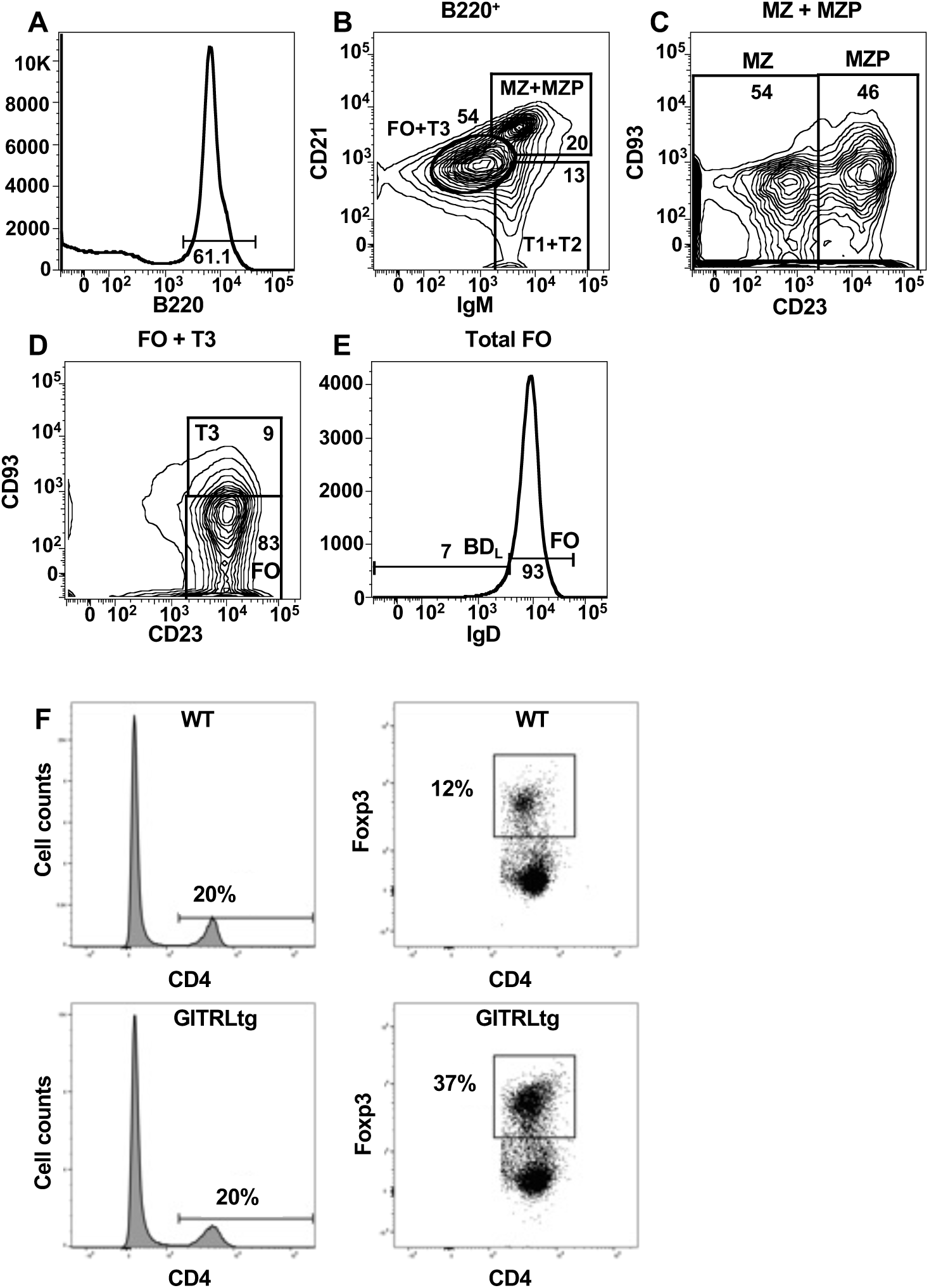
Splenic B cell subset and Treg flow cytometry gating strategies. Splenocytes were stained with B220, IgM, CD21, CD23, CD93 and IgD. B220^+^ B cells were gated (A) and analyzed for IgM and CD21 and IgM^+^CD21^+^ FO + T3 B cells or IgM^hi^CD21^+^ MZP + MZ B cells were gated (B). MZ and MZP were differentiated by expression of CD23 (C) and T3 and FO were differentiated by expression of CD93 (D). The FO gate was subsequently analyzed for IgD and BD_L_ were gated as IgD^low/-^ (D). Splenocytes were stained for CD4 and Foxp4 with CD4^+^ splenic T cells in WT and GITRLtg mice analyzed for Foxp3 expression to identify Treg (F).

**Supplemental Figure 2.**
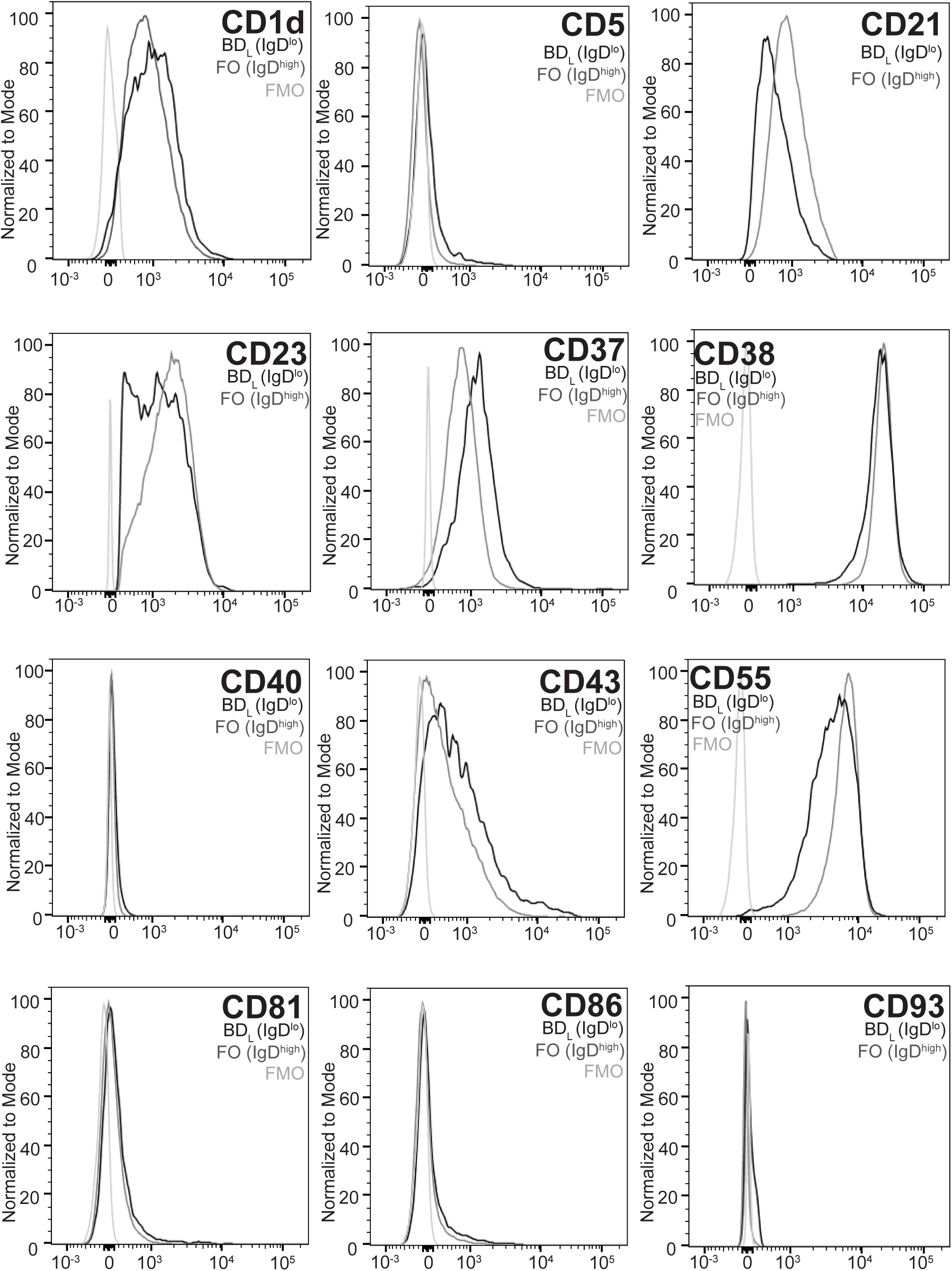
Cell surface expression of candidate BDL differentiating proteins. Splenocytes were stained for BD_L_ as for Suppl. Fig. 2 with the addition of an antibody specific to mouse CD1d, CD5, CD37, CD38, CD40, CD33, CD55, CD81, or CD86. BD_L_ (medium grey line) and FO IgD^hi^ (dark grey) B cells were gated as for Suppl. Fig. 1 and flow cytometry overlay histograms showing expression of CD1d, CD5, CD21, CD23, CD37, CD38, CD40, CD33, CD55, CD81, CD86, and CD93 are shown. The fluorescence minus one (FMO) control is shown in the light grey line.

## References

1. Baecher-Allan C, Kaskow BJ, and Weiner HL. Multiple Sclerosis: Mechanisms and Immunotherapy. Neuron. 2018;97(4):742–68.

2. Travers BS, Tsang BK, and Barton JL. Multiple sclerosis: Diagnosis, disease-modifying therapy and prognosis. Aust J Gen Pract. 2022;51(4):199–206.

3. Bluestone JA, McKenzie BS, Beilke J, and Ramsdell F. Opportunities for Treg cell therapy for the treatment of human disease. Front Immunol. 2023;14:1166135.

4. Wildin RS, and Freitas A. IPEX and FOXP3: clinical and research perspectives. J Autoimmun. 2005;25 Suppl:56–62.

5. Huter EN, Punkosdy GA, Glass DD, Cheng LI, Ward JM, and Shevach EM. TGF-beta-induced Foxp3+ regulatory T cells rescue scurfy mice. Eur J Immunol. 2008;38(7):1814–21.

6. O’Connor RA, and Anderton SM. Foxp3+ regulatory T cells in the control of experimental CNS autoimmune disease. J Neuroimmunol. 2008;193(1-2):1–11.

7. Stephens LA, Malpass KH, and Anderton SM. Curing CNS autoimmune disease with myelin-reactive Foxp3+ Treg. Eur J Immunol. 2009;39(4):1108–17.

8. Dawson NAJ, Vent-Schmidt J, and Levings MK. Engineered Tolerance: Tailoring Development, Function, and Antigen-Specificity of Regulatory T Cells. Front Immunol. 2017;8:1460.

9. Guo WW, Su XH, Wang MY, Han MZ, Feng XM, and Jiang EL. Regulatory T Cells in GVHD Therapy. Front Immunol. 2021;12:697854.

10. Goschl L, Scheinecker C, and Bonelli M. Treg cells in autoimmunity: from identification to Treg-based therapies. Semin Immunopathol. 2019;41(3):301–14.

11. Romano M, Fanelli G, Albany CJ, Giganti G, and Lombardi G. Past, Present, and Future of Regulatory T Cell Therapy in Transplantation and Autoimmunity. Front Immunol. 2019;10:43.

12. Ray A, Khalil MI, Pulakanti KL, Burns RT, Gurski CJ, Basu S, et al. Mature IgD(low/-) B cells maintain tolerance by promoting regulatory T cell homeostasis. Nat Commun. 2019;10(1):190.

13. Ray A, Basu S, Williams CB, Salzman NH, and Dittel BN. A novel IL-10-independent regulatory role for B cells in suppressing autoimmunity by maintenance of regulatory T cells via GITR ligand. J Immunol. 2012;188(7):3188–98.

14. Wolf SD, Dittel BN, Hardardottir F, and Janeway CA, Jr. Experimental autoimmune encephalomyelitis induction in genetically B cell-deficient mice. J Exp Med. 1996;184(6):2271–8.

15. Khalil MI, Gurski CJ, Dittel LJ, Neu SD, and Dittel BN. Discovery and Function of B-Cell IgD Low (BD(L)) B Cells in Immune Tolerance. J Mol Biol. 2021;433(1):166584.

16. van Olffen RW, Koning N, van Gisbergen KP, Wensveen FM, Hoek RM, Boon L, et al. GITR triggering induces expansion of both effector and regulatory CD4+ T cells in vivo. J Immunol. 2009;182(12):7490–500.

17. Dittel BN, Merchant RM, and Janeway CA, Jr. Evidence for Fas-dependent and Fas-independent mechanisms in the pathogenesis of experimental autoimmune encephalomyelitis. J Immunol. 1999;162(11):6392–400.

18. Allman D, Lindsley RC, DeMuth W, Rudd K, Shinton SA, and Hardy RR. Resolution of three nonproliferative immature splenic B cell subsets reveals multiple selection points during peripheral B cell maturation. J Immunol. 2001;167(12):6834–40.

19. Allman DM, Ferguson SE, Lentz VM, and Cancro MP. Peripheral B cell maturation. II. Heat-stable antigen(hi) splenic B cells are an immature developmental intermediate in the production of long-lived marrow-derived B cells. J Immunol. 1993;151(9):4431–44.

20. Lindsley RC, Thomas M, Srivastava B, and Allman D. Generation of peripheral B cells occurs via two spatially and temporally distinct pathways. Blood. 2007;109(6):2521–8.

21. Loder F, Mutschler B, Ray RJ, Paige CJ, Sideras P, Torres R, et al. B cell development in the spleen takes place in discrete steps and is determined by the quality of B cell receptor-derived signals. J Exp Med. 1999;190(1):75–89.

22. Allman DM, Ferguson SE, and Cancro MP. Peripheral B cell maturation. I. Immature peripheral B cells in adults are heat-stable antigenhi and exhibit unique signaling characteristics. J Immunol. 1992;149(8):2533–40.

23. Lightman SM, Utley A, and Lee KP. Survival of Long-Lived Plasma Cells (LLPC): Piecing Together the Puzzle. Front Immunol. 2019;10:965.

24. Page A, Hubert J, Fusil F, and Cosset FL. Exploiting B Cell Transfer for Cancer Therapy: Engineered B Cells to Eradicate Tumors. Int J Mol Sci. 2021;22(18).

25. Kubuschok B, Schmits R, Hartmann F, Cochlovius C, Breit R, Konig J, et al. Use of spontaneous Epstein-Barr virus-lymphoblastoid cell lines genetically modified to express tumor antigen as cancer vaccines: mutated p21 ras oncogene in pancreatic carcinoma as a model. Hum Gene Ther. 2002;13(7):815–27.

26. Winkler J, Tittlbach H, Schneider A, Vasova I, Strobel J, Herold S, et al. Adoptive transfer of donor B lymphocytes: a phase 1/2a study for patients after allogeneic stem cell transplantation. Blood Adv. 2024;8(10):2373–83.

27. Jeske AM, Boucher P, Curiel DT, and Voss JE. Vector Strategies to Actualize B Cell-Based Gene Therapies. J Immunol. 2021;207(3):755–64.

28. Cai Y, Schroeder JA, Jing W, Gurski C, Williams CB, Wang S, et al. Targeting transmembrane-domain-less MOG expression to platelets prevents disease development in experimental autoimmune encephalomyelitis. Front Immunol. 2022;13:1029356.

29. Duffy SS, Keating BA, and Moalem-Taylor G. Adoptive Transfer of Regulatory T Cells as a Promising Immunotherapy for the Treatment of Multiple Sclerosis. Front Neurosci. 2019;13:1107.

30. Bluestone JA, Buckner JH, Fitch M, Gitelman SE, Gupta S, Hellerstein MK, et al. Type 1 diabetes immunotherapy using polyclonal regulatory T cells. Sci Transl Med. 2015;7(315):315ra189.

31. Hefazi M, Bolivar-Wagers S, and Blazar BR. Regulatory T Cell Therapy of Graft-versus-Host Disease: Advances and Challenges. Int J Mol Sci. 2021;22(18).

32. Trzonkowski P, Bieniaszewska M, Juscinska J, Dobyszuk A, Krzystyniak A, Marek N, et al. First-in-man clinical results of the treatment of patients with graft versus host disease with human ex vivo expanded CD4+CD25+CD127-T regulatory cells. Clin Immunol. 2009;133(1):22–6.

33. Beres A, Komorowski R, Mihara M, and Drobyski WR. Instability of Foxp3 expression limits the ability of induced regulatory T cells to mitigate graft versus host disease. Clin Cancer Res. 2011;17(12):3969–83.

34. Kanamori M, Nakatsukasa H, Okada M, Lu Q, and Yoshimura A. Induced Regulatory T Cells: Their Development, Stability, and Applications. Trends Immunol. 2016;37(11):803–11.

35. Efe O, Gassen RB, Morena L, Ganchiku Y, Al Jurdi A, Lape IT, et al. A humanized IL-2 mutein expands Tregs and prolongs transplant survival in preclinical models. J Clin Invest. 2024;134(5).

36. Volfson-Sedletsky V, Jones At, Hernandez-Escalante J, and Dooms H. Emerging Therapeutic Strategies to Restore Regulatory T Cell Control of Islet Autoimmunity in Type 1 Diabetes. Front Immunol. 2021;12:635767.

37. Garces S, and Demengeot J. The Immunogenicity of Biologic Therapies. Curr Probl Dermatol. 2018;53:37–48.

38. Barnes RA, and Stallard N. Severe infections after bone marrow transplantation. Curr Opin Crit Care. 2001;7(5):362–6.

39. Bhatia S, Dai C, Landier W, Hageman L, Wu J, Schlichting E, et al. Trends in Late Mortality and Life Expectancy After Autologous Blood or Marrow Transplantation Over Three Decades: A BMTSS Report. J Clin Oncol. 2022;40(18):1991–2003.

40. Boffa G, Massacesi L, Inglese M, Mariottini A, Capobianco M, Moiola L, et al. Long-term Clinical Outcomes of Hematopoietic Stem Cell Transplantation in Multiple Sclerosis. Neurol. 2021;96(8):e1215–e26.

41. Rush CA, Atkins HL, and Freedman MS. Autologous Hematopoietic Stem Cell Transplantation in the Treatment of Multiple Sclerosis. Cold Spring Harb Perspect Med. 2019;9(3).

42. Msheik A, Assi F, Hamed F, Jibbawi A, Nakhl AM, Khoury A, et al. Stem Cell Transplantation for Multiple Sclerosis: A 2023 Review of Published Studies. Cureus. 2023;15(10):e47972.

43. Darrasse-Jeze G, Deroubaix S, Mouquet H, Victora GD, Eisenreich T, Yao KH, et al. Feedback control of regulatory T cell homeostasis by dendritic cells in vivo. J Exp Med. 2009;206(9):1853–62.

44. Bar-On L, Birnberg T, Kim KW, and Jung S. Dendritic cell-restricted CD80/86 deficiency results in peripheral regulatory T-cell reduction but is not associated with lymphocyte hyperactivation. Eur J Immunol. 2011;41(2):291–8.

45. Mann MK, Maresz K, Shriver LP, Tan Y, and Dittel BN. B cell regulation of CD4+CD25+ T regulatory cells and IL-10 via B7 is essential for recovery from experimental autoimmune encephalomyelitis. J Immunol. 2007;178(6):3447–56.

46. Stuart T, Butler A, Hoffman P, Hafemeister C, Papalexi E, Mauck WM, 3rd, et al. Comprehensive Integration of Single-Cell Data. Cell. 2019;177(7):1888–902 e21.

47. Aran D, Looney AP, Liu L, Wu E, Fong V, Hsu A, et al. Reference-based analysis of lung single-cell sequencing reveals a transitional profibrotic macrophage. Nat Immunol. 2019;20(2):163–72.

48. Immunological Genome P. ImmGen at 15. Nat Immunol. 2020;21(7):700–3.

49. Waltman LvE, N.J. A smart local moving algorithm for large-scale modularity-based community detection. Eur Phy J B. 2013;86.

50. Becht E, McInnes L, Healy J, Dutertre CA, Kwok IWH, Ng LG, et al. Dimensionality reduction for visualizing single-cell data using UMAP. Nat Biotechnol. 2018.

51. Finak G, McDavid A, Yajima M, Deng J, Gersuk V, Shalek AK, et al. MAST: a flexible statistical framework for assessing transcriptional changes and characterizing heterogeneity in single-cell RNA sequencing data. Genome Biol. 2015;16:278.

52. Cao J, Spielmann M, Qiu X, Huang X, Ibrahim DM, Hill AJ, et al. The single-cell transcriptional landscape of mammalian organogenesis. Nature. 2019;566(7745):496–502.

53. Holland CH, Szalai B, and Saez-Rodriguez J. Transfer of regulatory knowledge from human to mouse for functional genomics analysis. Biochim Biophys Acta Gene Regul Mech. 2020;1863(6):194431.

